# Moving as a group imposes constraints on the energetic efficiency of movement

**DOI:** 10.1101/2023.11.26.568763

**Authors:** James A. Klarevas-Irby, Brendah Nyaguthii, Damien R. Farine

## Abstract

Movement is a key part of life for both solitary and group-living animals. In solitary animals, the energetic costs of making large displacements can be mitigated by energetically efficient strategies—specifically faster, straighter movements. However, little is known about whether moving as part of a collective enhances or limits the ability for individual group members to express such strategies. Drawing on 6 years of population-level high-resolution (1Hz) GPS tracking of group-living vulturine guineafowl (*Acryllium vulturinum*), we detected 886 events from 94 tagged individuals where their groups made large displacements, shifting their home ranges in response to changing environmental conditions. We contrasted these movements with data with 94 large displacement events by 19 lone, dispersing individuals. Our results suggest that individuals moving as part of group can significantly reduce their energetic cost of transport when making large displacements (15.5% reduction relative to their normal daily ranging behaviours) by increasing the speed and straightness of their movements. However, even during their most-efficient movements, individuals in groups could not achieve or maintain the same increases in speed that others could when moving alone, resulting in significantly limited efficiency gains (solitary individuals were 34.9% more efficient than individuals in groups). Overall, this study provides evidence for a substantial, and previously hidden, energetic cost arising from collective movement.

## INTRODUCTION

Individuals in groups have to solve the same challenges as solitary individuals, from finding resources and mates [1,2] to escaping from predators[3,4]. While individuals in groups can gain some advantages over individuals in terms of detecting predators [5], improved navigation [6,7], and collective memory [8], they also have to overcome the challenges of coordinating their actions to remain cohesive. Disagreements among individuals in terms of their preferred movement directions [9] or movement capabilities [10] can require individuals to compromise their own optimal strategy in order for the group to maintain cohesion [11]. Such compromises are likely to impose costs on group members, but to date the nature and extent of these cost remain largely overlooked. One potential cost that individuals in collectives could pay is through an increased cost of transport—i.e. the energetic cost incurred to move a given distance. In many species, the energetic costs incurred through movements have led to the evolution of strategies that can increase energetic efficiency when making large displacement. For example, migrating [12–14] and dispersing [15–19] animals typically move faster and straighter, allowing them to conserve energy over large distances [20]. A key question is whether groups can achieve similar increases in efficiency, relative to individuals, when making movements over large distances or whether the challenges of coordinating actions as a group limits the ability for individuals to express energetically efficient movement strategies.

Allometric models of the energetic costs of movement [21,22] reveal that higher movement speeds require increased energetic investments, but result in an overall reduction in the cost of transport. This relationship between movement speed and efficiency in the cost of transport can help explain some patterns in terms of how groups move, for example in heterogenous groups where differences in body size correspond to differences in movement capability. In olive baboons (*Papio anubis*), smaller individuals have a slower natural gait than larger individuals, but group speed is determined by the larger individuals, with smaller group members having to speed up [10]. This pattern makes sense in the context of the cost of transport, as while smaller individuals increase their instantaneous energy output, they ultimately increase their efficiency, whereas if larger individuals slowed down to the speed of the slowest individuals they would incur a higher cost of transport. Thus, compromise in animal groups may entail an increase in the instantaneous costs paid by some individuals, but ultimately increase the overall efficiency of group members’ movements. Other forms of compromise, such as making collective decisions about where to go next (e.g. through shared decision making [9]) could also impact movement costs. For example, an individual initiating movement in a desired direction and failing requires a net output of energy with no net displacement achieved, thereby reducing efficiency. Further, while conflicts are being resolved most or all of the group members may have to slow down or stop entirely. The continuous process of decision-making ‘on the go’ [23] could reduce the speed and the continuity of movement of groups. As a consequence, we expect that moving as part of a group could result in less efficient movements for individuals compared to moving alone, potentially resulting in a greater cost of transport.

While we predict that moving in groups should impose some constraints on efficiency, it is important to consider that the net costs of displacement are determined not only by the speed of movement but also by the straightness of the path taken between two points. More tortuous paths, regardless of speed, involve covering more ground and thus spending more total time moving than straighter paths—thus inducing a net energy cost. When moving in a group, pooling directional uncertainty across multiple individuals can yield greater navigational accuracy (the ‘many wrongs’ hypothesis [7,24]). Studies in pigeons (*Columba livia*) [6,25,26] and humans [27] have confirmed that individuals are more accurate at navigating towards targets when moving in a group than when moving alone, resulting in more-directed overall movement paths. Thus, individuals may benefit from living in groups—and potentially offset slower movements—if the collective navigational capacity of groups allows them to follow straighter movement paths.

In this paper, we combine high-resolution GPS tracking of vulturine guineafowl (*Acryllium vulturinum*), an almost exclusively terrestrial bird, with laboratory models of guineafowl physiology [28] to compare the cost of transport for individuals moving in groups versus alone. Specifically, we use our data to generate a two-part comparison of the movement characteristics of] individuals moving as part of a collective (Figure 1A). First, we compare the normal daily movements of individuals in groups to their movements on days when their group made large displacements. This first comparison allows us to test if individuals in groups are capable of increasing their energetic efficiency (i.e. through a lower cost of transport) under conditions that should favor the expression of more-efficient movements. Second, we then compare these data to similar large movements of individuals moving alone, captured during the solitary phase of dispersal. This second comparison allows us to determine the scale of any increases in efficiency expressed by individuals in groups versus lone individuals. Across these three conditions, we also analyze the properties of individuals’ movements, thus quantifying the relative contributions of movement speed, continuity, and straightness to their energetic efficiency. In doing so, we demonstrate that individuals pay greater energetic costs of transport during collective movements, arising from reductions in the speed and continuity of movement, revealing an often-overlooked cost of living in groups.

**Figure 1.**
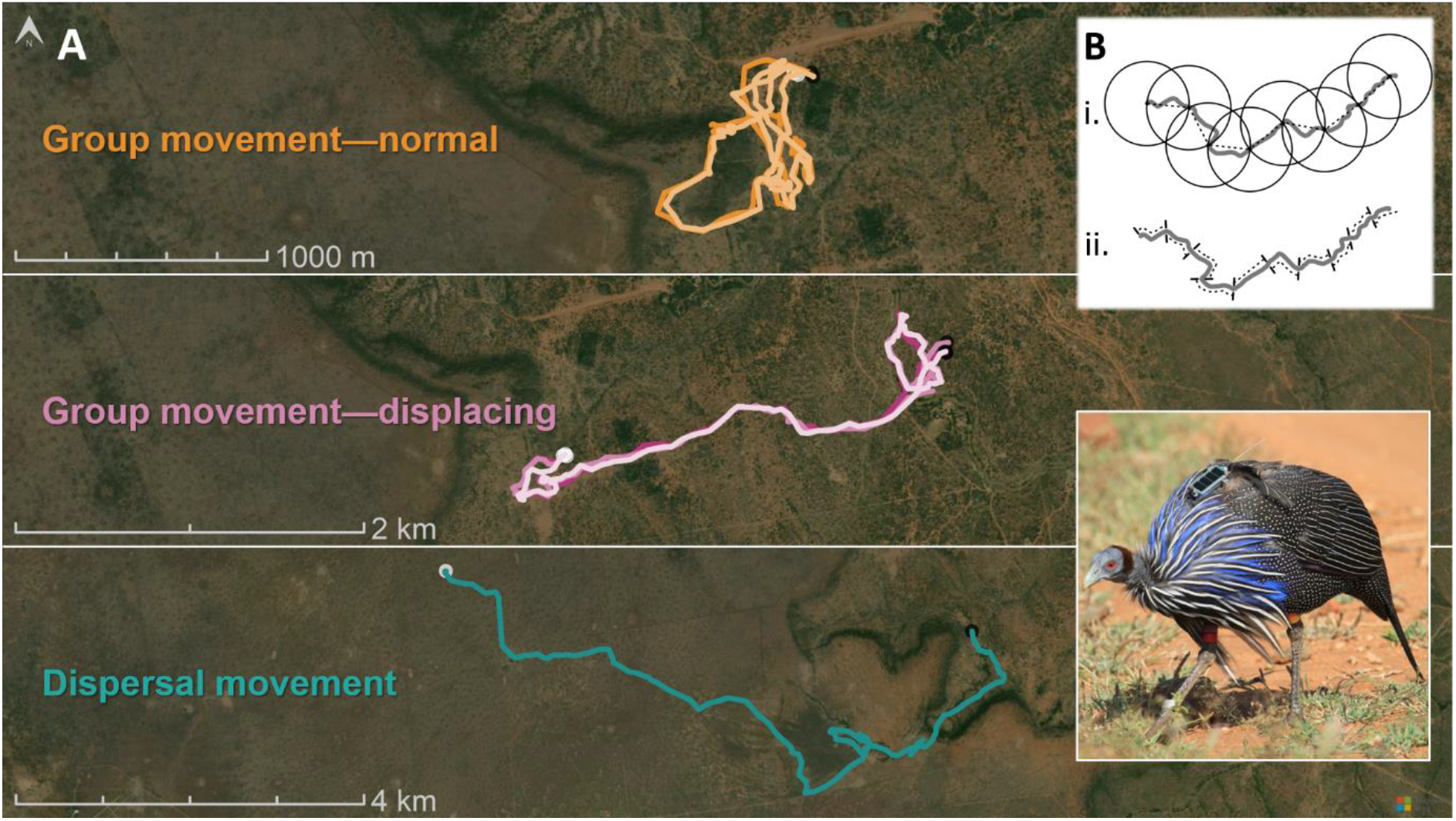
Movement tracks of GPS-tracked vulturine guineafowl across three conditions. (A) Illustrative tracks of normal daily movements of birds moving as part of a cohesive social group (orange, top), movements made by those same groups on days when they make large displacements in search of resources (purple, middle), and large displacements made by lone individuals during natal dispersal (green, bottom). Morning and evening roosting points marked by black and white points, respectively. (B) Illustration of how a track segmented into 50m units of either (i) net or (ii) cumulative displacements. Inset shows a vulturine guineafowl fitted with a solar-powered GPS tracker (photo: D. Farine).

## METHODS

### Study system

We conducted our research at the Mpala Research Centre in Laikipia, Kenya (0.292120, 36.898670), which comprises in a mix of semi-arid savannah and scrubland habitats. Here, we have studied a population of vulturine guineafowl (*Acryllium vulturinum*)— consisting of 600–1000 individuals across approximately 20 stable groups— continuously since 2016. Since the start of this study, over 1300 individuals have been identified and marked with individually-numbered stainless-steel rings, and a unique visual identifier—either a four-color combination of plastic leg bands, or a numbered canvas tag on the wing (for individuals captured as juveniles and too small to be fitted with rings). Birds were captured in groups, using large (8m x 4m x 2m) walk-in traps to catch all members of a group at once.

#### GPS tracking

A subset of individuals in each group were fit with a 15g Bird Solar GPS tag (e-obs GmbH), using a backpack style Teflon harness and a foam rubber pad to elevate solar panels above body feathers (following Papageorgiou et al., 2019). While the exact number of tagged individuals varied during the study period, there were typically between 2 and 5 tags deployed to residents in each social group (see He et. al., 2023 for details on deployment strategies), with an additional 25 tags deployed on subadult females to track dispersal, resulting in approximately 100 concurrently-deployed tags at any given time. In total, the approximate weight of all markings and GPS backpacks was 20.5g, less than 2% of birds’ body weight.

For the purpose of this study, we used GPS tracking data from 113 individuals, comprising 19 dispersing subadult females—approximately 18 months old at the time of dispersal— and 94 group-living residents (52 adult males, 42 adult females) distributed across all social groups in our study population. GPS data were recorded over 5 years from Nov. 2016 to Sep. 2021, including two periods of dispersal in April and October 2019.

GPS devices were programmed to record data during daylight hours, from 06:00 to 19:00, when birds were active outside of their nighttime roosts. GPS data (date, time, and location) were recorded at two resolutions, depending on tag battery level. When tags’ batteries were fully charged (approximately every second to third day), tags collected continuous 1Hz data (i.e. one fix per second). When battery levels fell below the high-resolution charge threshold, we set tags to record a 10 second burst of 10 fixes every five minutes. If battery charge was at the lowest threshold, tags were set to record one point every 15 minutes (this threshold was not crossed during this study). Data were remotely downloaded every two days (sometimes daily in the case of actively-dispersing birds) using a BaseStation II (e-obs Digital Telemetry, Grünwald, Germany). Data were separated into two resolutions for analytical purposes: high-resolution data, comprising all continuous periods of 1Hz data; and five-minute data, comprising data from the tenth second of every fifth minute of the day. The latter was collected from both the low-resolution dataset and by sub-sampling the 1Hz data, thereby reliably providing one fix every 5 minutes for every bird on every day of tracking. Accelerometer functions of GPS tags was disabled for this study, as the increased amount of data would have hindered our ability to reliably download data at regular intervals, given the number of tags concurrently deployed. GPS Data were uploaded to Movebank (https://www.movebank.org/) and retrieved and prepared for analysis in R using the move package [31].

### Research Permits

All work was conducted under research permits from the Max Planck Society Ethikrat Committee (2016_13/1), the National Commission for Science, Technology and Innovation of Kenya (NACOSTI/P/16/3706/6465; NACOSTI/P/21/8699), the National Environment Management Authority (NEMA/AGR/68/2017), under a Research Authorisation and a Capture Permit issued by the Kenyan Wildlife Service, research authorisations from the Wildlife Research and Training Institute, and in affiliation with the National Museums of Kenya.

### Analysis

All analyses were performed in R version 4.0 [32].

#### Identifying large-scale movements

In order to draw comparisons between movements of grouped individuals and lone dispersers, we isolated all days within our dataset of GPS movements that comprised of a large daily displacement. On most days, groups of guineafowl return to the same roost, or to a nearby roost, resulting in small daily displacements. However, on the onset or end of extreme weather conditions (e.g. droughts), groups move out of (or back into) their regular home range [33]. During such days, groups make similar displacements to actively-dispersing individuals, and through similar habitats as they navigate their way outside of the core population’s range. Following [34], we identified days which primarily contained large-scale movements based on the length and straightness of each individual’s daily movement path—if either the roost-to-roost distance (i.e. daily net displacement) was greater than 1500m, or if the roost distance was greater than 1200m and the ratio of the distance between roosts and their total daily track length was greater than 0.3. The latter captures days of large movements where groups or individuals made a large turn, thereby reducing the roost-to-roost distance but maintaining a large overall movement distance. These same criteria were used for isolating the solitary movements of dispersers, as transience in this species typically entails two modes: active dispersal, where individuals make large solitary movements, and more local days of movement which observations suggest correspond to periods of movement within groups other than their natal or post-settlement group [34].

For grouped individuals, we also included a matched set of “normal” days of movement to test for changes in the cost of transport (and subsequent efficiency benefits) on days of extreme movements. For each individual, we selected N days at random from the remaining data set, where N was the number of days of large movements recorded from that individual. For lone dispersers, we only used days of extreme movement to facilitate comparison with grouped birds, as overall changes in movement during dispersal were characterized in Klarevas-Irby, Wikelski, & Farine, 2021.

#### Defining movement states

We employed an unsupervised Hidden Markov Model (HMM) to identify the different states of movement exhibited by vulturine guineafowl. Movement states were characterized by first summing the distance moved and absolute turning angles for every 10 seconds in the high-resolution data, a 1Hz resolution would otherwise violate the Markov assumption. We separated movement into four states across the entire high-resolution dataset, using the R package depmixS4 [35], with our three categorical labels (grouped individuals making extreme movements, grouped individuals moving normally, and solitary dispersers) fitted as a covariate on the state transition matrix. The large-scale movements of grouped individuals were treated as the reference category when fitting state transition probabilities. The choice to use a four-state model was based on field observations that individuals spend time not moving (state 1), making slow, tortuous foraging movements (state 2), walking at a medium speed (state 3), and moving quickly in a directed manner (state 4). A critical reason for the implementation of 4-state model was the need to isolate a clear “stationary” state for the purposes of calculating metabolic expenditure—birds were considered to be moving when assigned to states 2-4. Once states were assigned for each 10-second cluster of data, we then attributed each given state to all 1Hz data points which contributed to each cluster (see Supplementary Table S1 for state distributions and transition probabilities).

#### Characterizing movement behaviours

We extracted three representative measures to characterize birds’ movement behaviours: daily track length (km), speed while moving (m s^-1^), and the straightness of movement (a straightness index, [36]). To control for variation in high-resolution data collected across different days, daily track lengths were measured from the sum of displacements in the five-minute data. Movement speeds were calculated as the speed-while-moving (i.e. from all data assigned to states 2-4) based on the per-second displacement, summarized for each 5-minute window of available high-resolution data. To reduce the effect of small GPS errors on calculated velocities, we derived individuals’ speeds at each second from the mean velocity over a rolling 5-second window within the high-resolution data (i.e. for each second of 1Hz data, we averaged the individual’s speed with the two seconds which preceded and followed it). Straightness of movement was characterized by dividing the net displacement achieved over each available 5-minute window of 1Hz data by the summed cumulative displacement therein.

To test how the behaviours of individuals in groups varied during periods of extreme movement, we first fit linear mixed models (LMM), using the package lmerTest [37], to each movement measure. When analyzing daily track length, we considered one measure per individual per day. Because the distribution of daily track lengths is left-truncated, forming a long-tailed distribution, we log-transformed values to aid with model fitting. For analyses of movement speed and straightness, we included one observation from each available 5-minute period of continuous high-resolution data (mean=69.4, range=4 to 156 observations per individual per day). In each model, we included context (i.e. group member during a large displacement, group member on a normal day, or lone disperser) as a predictor, and individual identity as a random intercept. All models were fit using restricted maximum likelihoods (REML). Specific equations for all LMMs can be found in the corresponding table in the supporting information (Supplementary Tables S2-S4).

#### Calculating the energetic cost of movement

To quantify the energetic costs of movement, we used published data [28] on the relationship between metabolic costs (mL O_2_ kg^-1^ s^-1^) and movement speed (m s^-1^) in the closely-related, and morphologically-similar, helmeted guineafowl (*Numida meleagris*). We used two separate formulas to calculate the costs incurred for each second of high-resolution data, depending on whether birds were either stationary (state 1) or moving (states 2–4). When moving, guineafowl exhibit a linear relationship between speed (i.e. velocity v > 0) and oxygen consumption, given by VO_2_ = (24.0v + 27.2), where V0_2_ is the per-minute volume of oxygen consumed in mL O_2_ per kilogram of body mass (mL O_2_ kg^-1^ min^-1^). The formula for when birds were stationary is a fixed consumption of 19.1 mL O_2_ kg^-1^ min^-1^, corresponding to the oxygen consumption rate when not moving (i.e. the resting metabolic rate), as described in the literature [28]. We then transformed the per-second measures of metabolic oxygen consumption into units of Joules kg^-1^ s^-1^ using a conversion factor of 20.1 J mL^-1^ O_2_ (per Ellerby et al. 2003; Marsh et al. 2006).

#### Calculating the energetic cost of transport

To relate energetic expenditure to achieved displacements (i.e. the cost of transport), we partitioned movement tracks in our high-resolution data into fixed segments representing 50 meters of net or cumulative displacement (Figure 1B). Net displacement is the absolute movement in space (i.e. a straight-line distance) between two points in time, while cumulative displacement is the sum travel distance of all individual steps. We defined net displacement segments starting from the first available second of high-resolution data (i.e. after the GPS switched on or switched from low-resolution to high-resolution) until the 50 m net displacement threshold was crossed. We then used the first GPS point to fall on or outside of the 50 m radius as the first point for the following segment. We also calculated the corresponding cumulative displacements from the same high-resolution data by summing each consecutive step length in a track until it reached a sum of 50 m, at which point we started a new segment. We then summed the per-second-costs for each detection which contributed to a segment to calculate the total energetic expenditure for that segment, and translated it into the energetic cost of transport (in J kg^-1^ m^-1^) for each type of displacement by dividing by the distance travelled (either net or cumulative). Because segments varied slightly from perfect, 50 m denominations, cost of transport values were calculated relative to the true distances travelled.

For each displacement type, we fit separate LMMs with the cost of transport associated with each segment as the response variable, context as a predictor, and individual identity as a random intercept. Model formulations can be found in corresponding supplementary tables (Supplementary Tables S5-S6).

#### Calculating cumulative daily energy use

We calculated the total energetic expenditure over each 13-hour day for each bird by summing all of the per-second-costs from each unique day of tracking for each individual into a single measure of daily energetic expenditure (i.e. J kg^-1^ day^-1^). Not all days contained an equal amount of high-resolution data (average of 5.75 hours of high-resolution data on days when high-resolution data were recorded), and thus we standardized the cost value for each day by multiplying the mean per-second-cost within the day by 46800 seconds. Days containing fewer than 2 hours of high-resolution data were excluded from this analysis, to avoid potentially over-representing days with too little high-resolution data.

To estimate the change in energy use across contexts, we fit an LMM of daily energetic expenditure as the response variable, with context as the predictor variable and individual identity as a random intercept. Model formulation can be found in Supplementary Tables S2-S7.

## RESULTS

### Individuals moving in groups increase their energetic efficiency when making large displacements

We extracted 886 days of large daily displacements from population-scale GPS tracking starting in September 2016 until September 2021. When moving as part of a group, individuals were more energetically efficient on days when the group made a large displacement (average 15.5% reduction in the net cost of transport, p=0.03, 8.4% reduction in cumulative cost of transport, p<0.001, Figure 2, Tables S5-S6). On average, individuals moving in groups expressed a 17.3% increase in daily travel distances on days of large movements (p<0.001, Figure 3A, Table S2) relative to their normal movements, which resulted in a 2.9% increase in total daily energy expenditure (p<0.001, Figure S1, Table S3).

**Figure 2.**
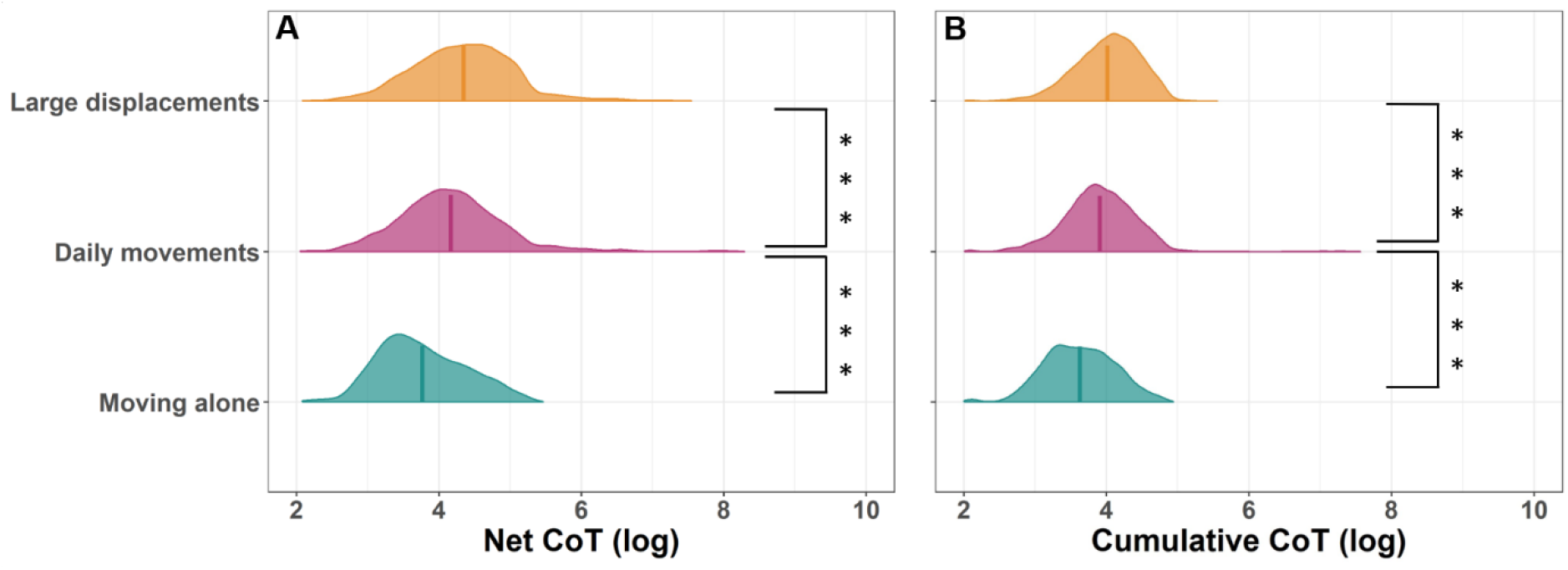
The cost of transport (CoT) is significantly reduced when individuals make large displacements relative to regular daily movements, but individuals moving in groups are less efficient than those moving alone. Plots show log-transformed distributions of expressed costs of transport for individuals making (A) a 50 m net displacement and (B) a corresponding cumulative displacement to achieve 50 m of net displacement. Vertical lines show the mean cost of transport values per category, statistical significance in differences between categories marked with stars (*** p<0.001, see Tables S4-S5 for full model results).

**Figure 3.**
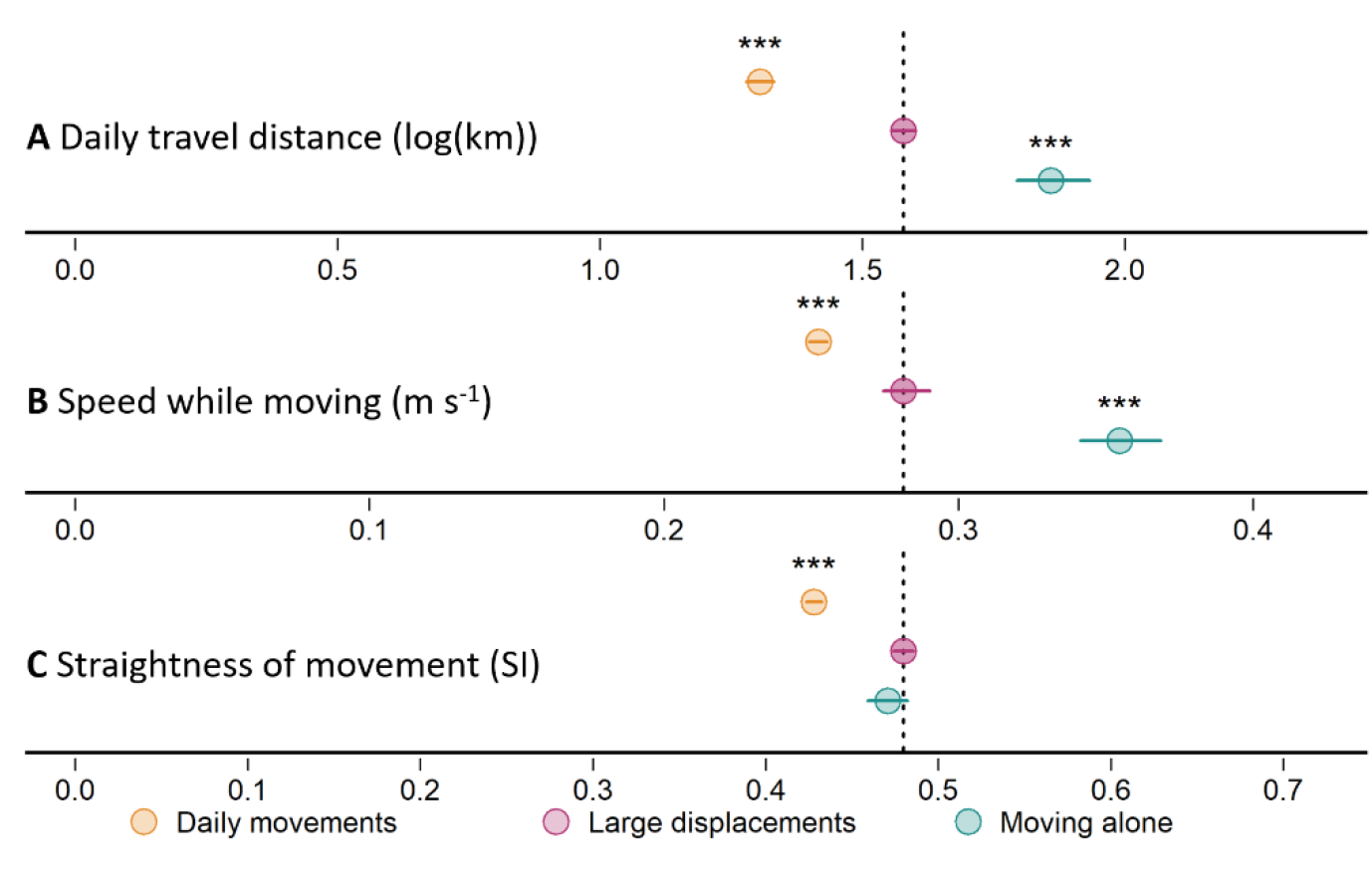
Birds moving in groups moved farther, faster, and straighter when making large displacements, but lone individuals moved even further and faster. (A-C) summaries of linear mixed-effect models (LMMs) characterizing (coefficient±95% confidence intervals) movement behaviours when individuals made daily movements with their group, made large displacements in a group, and moved alone during dispersal. Significance (*** p<0.001) was estimated using large displacements of individuals in groups as the reference category (dotted vertical lines correspond to model intercept). Full model results available in Tables S2, S6, S7.

### Individuals moving in a group are less efficient than individuals moving alone

Comparing 96 days of large dispersal movements by individual moving alone to the large movements of individuals moving in groups shows that individuals moving in groups were significantly less efficient than lone dispersers (average 32.3% greater net cost of transport, p<0.001, 25.1% greater cumulative cost of transport, p<0.001, Figure 2, Tables S4-S5). Lone dispersers travelled on average an additional 17.8% further per day than individuals when making large movements in groups (p<0.001, Figure 3A, Table S2), expending only 5.6% more total energy per day (p<0.001, Figure S1, Table S3).

### Individuals moving in groups make slower, more sporadic movements

An analysis of fine-scale movement behaviours revealed that individuals moving in groups expressed distinct differences in behaviour, relative to individuals moving alone, when making large displacements. The increased energetic efficiency of group members’ large movements was the result of a significant increase in average speed-while-moving (10.2 %, p<0.001, Figure 3B, Table S6), and a significant increase in the average straightness of movement (10.8%, p<0.001, Figure 3C, Table S7). However, lone individuals were significantly faster on average (26.1% greater, p<0.001, Figure 3B, Table S6), with no clear difference in the straightness of movement (1.9% lower than group members, p=0.77, Figure 3C, Table S7) relative to individuals making large displacements as part of a group.

Categorical differences in state transition probabilities within our HMM revealed that individuals moving in groups also had a substantially lower likelihood of maintaining the most efficient movement state (65.2% probability of remaining in state 4, which corresponds to the fastest, straightest movements, Table S1) than individuals moving alone (75.2% probability of remaining in state 4, Table S1). In general, group members showed a greater probability of transitioning from “slower” to “faster” states (e.g., transitioning out of the stationary first state, or transitioning to states 3 and 4 from state 2) during days of large movement than they did during normal daily movements (Table S1), but this general probability of transitioning to higher states was greater for lone dispersers.

## DISCUSSION

Our study shows that group living can have a significant, constraining effect on the energetic efficiency of movement for individuals that move as part of a group. While individuals are more efficient when making large displacements in a group relative to their typical daily ranging movements in the same groups, during large displacements they do not move as fast or as continuously as individuals that make similarly large displacements alone. This means that individuals that move alone can achieve over 35% larger travel distances while only using slightly less energy for movement than individuals moving as part of a group. These results reveal that group living is likely to generate greater mobility costs for individuals.

While many studies on collective movement have focused on the differences in behaviour expressed by groups of different sizes [39,40], few have compared individuals in groups versus solitary individuals. In part, this is due to the inherent challenges in trying to study group-living animals making ecologically relevant movements without their group. Here, we were able to leverage a dataset of spanning multiple life history stages to draw direct comparisons between the behaviours of individuals moving in groups versus dispersers that moved alone. While these movements represented distinct life stages (being in a group versus dispersing), the two sets of movement do overlap. Vulturine guineafowl exhibit delayed dispersal (in our dataset, between 1.5 and 2.5 years post-hatching), meaning that their movements are made after they have gained substantial life experience and generally reached adult body size (females reach 90% of their adult body size at 1 year of age, *D.R. Farine, unpublished data*). Further, groups will make large displacements even when they contain females that have not yet dispersed, despite limitations to their daily displacements when groups have young chicks [39]. Thus, we expect that the observed differences do not reflect changes associated with physiology or life stages *per se*.

Our results raise the question: what aspects of movement behaviour can individuals optimize when alone and which are limited when in a group? A previous study on the same species [20] found that dispersing individuals increase the speed and straightness of their movements by 27.7% and 10.3%, respectively, when compared to their normal non-dispersing movements. Studies in other systems [17–19] reported similar increases in both movement speed and straightness (and greater increases in speed than straightness)—such as a 23.9% increase in speed compared to a 10.3% increase in straightness in dispersing lions (*Panthera leo)* [16]. In vulturine guineafowl, individuals moving in groups exhibited a nearly-identical increase in path straightness as individuals moving alone, but achieved a substantially lower increase in speed (10.2%) relative to their regular daily movements. These results suggest that speed is the main limiting factor for the efficiency of individuals in groups, more so than the straightness of their paths. Previous work on baboons has demonstrated how preferred movement speed within a group is in conflict among group members [10], with individuals either having to move faster when trailing behind, or else stopping-and-starting if they outpaced the group. These patterns have consequences for energy efficiency. While slower individuals increase their speed (and thus overall efficiency), mechanical (e.g., limb length) disadvantages mean that they are likely to experience diminishing returns in energetic efficiency as they move beyond an efficient gait. At the other end of the spectrum, faster individuals have to repeatedly pause their movements, introducing substantial efficiency costs [41].

A reduction in the continuity of movement (i.e., pausing to wait for others) is likely to be exacerbated by the process of collective decision-making. Groups face significant coordination challenges, and most groups resolve these conflicts by making so-called shared decisions (i.e. voting for a direction) [9,11,42,43]. Previous work using GPS to track the movement of baboon troops found that individuals are less likely to move when, for example, there is a larger conflict over the proposed directions of travel [9,44]. We found evidence that corroborates this effect as playing an important role in driving our results. We found different state transition probabilities within our HMM based on whether individuals were making large displacements in a group or doing so alone. Specifically, we found that individuals in groups were less continuous in their movements, stopping and starting more frequently than solitary individuals (captured by a lower probability of remaining in the fast movement state). Our results highlight that collective movements, while beneficial in some respects, can also be costly in other respects.

Extensive work on collective navigation, focusing on the performance of lone individuals vs groups, especially using homing pigeons, has suggested that groups should gain navigational benefits that reduces the overall distance travelled [6,25,26,45]. These benefits are predicted to arise from the many-wrongs hypothesis, where the error in each individuals’ directional estimates cancel out to produce a more accurate average when integrated at the group level [7,24]. Our findings that individuals did not move straighter when making large displacements as a group relative to moving alone raises the question of whether vulturine guineafowl do not benefit from collective navigation. There are two points to consider, both of which highlight some interesting dynamics in terms of movement. First, in our study we considered straightness over a relatively small period (5 minutes of movement), a timeframe over which solitary dispersers appear to be able to match the straightness of individuals moving in groups. However, we observed many cases in which dispersers made large turns in the middle of a day, which would dramatically reduce the straightness when considered over the course of an entire days’ movement (or an entire displacement period). This leads to the second point, which is that solitary dispersers are less likely than groups to be moving in a goal-oriented way, because dispersers’ movements must include some level of exploration. While straight-line paths are thought to be some of the most effective for sampling for settleable habitat, this utility quickly breaks down under energetic constraints [46], and doesn’t account for the notion that naïve individuals may be more likely to encounter movement barriers [47]. As such, dispersers may face certain constraints to the straightness of their movements, with increasing movement speeds allowing them to increase their sampling area, in addition to the energetic benefits conferred. Groups, by contrast, are more likely to benefit from greater navigational efficiency [45], especially when movements are goal-oriented, over the course of an entire displacement event. Thus, while collective movements can impose significant limitations to the efficiency of movement speeds, group members may be able to reclaim some benefits over their solitary counterparts if the costs of transport are estimated in a goal-specific way (moving between habitats for individuals in groups versus settling in a new group for dispersing individuals).

A further factor that could modulate movement behaviours during large displacements is the differing habitat selection pressures acting on dispersers and residents. This again is a multidimensional topic. For example, several studies have shown that animals tend to select for different habitats during large movements than those they typically reside or forage in [48,49]. However, if animals are searching for foraging resources (as was likely the case for the large group movements in our study), then they may need to move through more-resistant habitats to do so. By contrast, dispersing guineafowl must locate other groups to enter. While this might also force them into residential habitats, the range at which a group can be detected (e.g., via auditory cues like contact calls) is possibly large enough to allow dispersers to focus on moving through the most suitable habitats without sacrificing sampling opportunities. Conversely, solitary individuals are also likely to be substantially more sensitive to predators [50,51], which could limit their access to certain habitats (e.g. avoiding very open or very dense vegetation)or restrict their activity at times when predators are most active [52]. Previous work on vulturine guineafowl has suggested that dispersers disproportionately select more open habitats, and that these provide substantial benefits in terms of movement speed [53]. Groups, by contrast, may be less restricted by predator activity [54], in both space and time, lending to a greater flexibility in potential movement behaviours, or may be more restricted to searching for resources in more vegetated habitats that do not promote efficient movements.

Despite the clear costs that individuals pay in terms of their energetic efficiency when moving as a group, group living could still bring substantial benefits in other ways. One major energetic cost for many vertebrates is brain power [55–57], and moving as a group could modify these costs. Examples of lone ants and pigeons being unable to recapitulate routes after following others [58,59], suggest that followers may be able to shift their cognitive resources from landscape features to focus on other factors, such as the behaviour of conspecifics or watching for predators. Given that cognitive load can be measured as birds navigate through landscapes [60], future studies could compare the cognitive costs paid by individuals when navigating large displacements alone versus in a group. Similarly, biologging advances in heart rate monitoring [61,62] could also potentially reveal whether individuals pay lower physiological costs (e.g. maintaining a lower heart rate) when moving in a group versus alone. There are several ways this could happen, one being because they are less sensitive to risk [63,64], and another because following can reduce the heart rate when moving (drafting during races is a classic example in humans, Bilodeau et al., 1994; Coast & Piatt, 2001). These complex interactions of risk [67,68], energetics [69,70], resources [2,71], and social structures [72] all provide fertile ground for future research into how group living modifies the fundamental ecology of social species.

## ACKNOWLEDGEMENTS

We thank the Mpala Research Centre, the Kenyan Wildlife Service, and the Ornithological Section of the National Museums of Kenya for supporting this research work. We are especially grateful to Dr. Peter Njoroge from the NMK for the long-term collaboration on our project, and Dr. Fred Omengo for his continued advice and oversight on behalf of the KWS. We also thank the Ol Jogi Wildlife Conservancy, Mr. Peter Jessel, and El Karama Ranch for their support in allowing us to track dispersing guineafowl on their properties. We are grateful to Wismer Cherono, John Wanjala, Monicah Wambui, John Ewoi, Martha Sisto, and Danai Papageorgiou for their contributions to data collection and field assistance.

---

**Supplementary Table S1.** Summary of the fitted Hidden Markov Model (HMM) across all movement data (i.e., including grouped and solitary movements) which extracted 4 states of movement based on step lengths and turning angles. State transition probabilities were fit to vary according to categorical label (i.e., local grouped movements, large grouped movements, or solitary dispersal).

| Summary of 4-state HMM of Vulturine Guineafowl Movement |  |  |  |  |
| --- | --- | --- | --- | --- |
| Initial State Probabilities |  |  |  |  |
| State 1 | State 2 | State 3 | State 4 |  |
| 0.127 | 0.385 | 0.245 | 0.243 |  |
| State Transition Probabilities |  |  |  |  |
| Large movements (grouped) |  |  |  |  |
|  | To State 1 | To State 2 | To State 3 | To State 4 |
| From State 1 | 0.762 | 0.037 | 0.184 | 0.017 |
| From State 2 | 0.043 | 0.937 | 0.018 | 0.002 |
| From State 3 | 0.267 | 0.007 | 0.557 | 0.168 |
| From State 4 | 0.043 | 0.001 | 0.304 | 0.652 |
| Local movements (grouped) |  |  |  |  |
|  | To State 1 | To State 2 | To State 3 | To State 4 |
| From State 1 | 0.796 | 0.037 | 0.157 | 0.009 |
| From State 2 | 0.047 | 0.936 | 0.015 | 0.001 |
| From State 3 | 0.269 | 0.006 | 0.597 | 0.127 |
| From State 4 | 0.042 | 0.002 | 0.334 | 0.622 |
| Large movements (solitary) |  |  |  |  |
|  | To State 1 | To State 2 | To State 3 | To State 4 |
| From State 1 | 0.699 | 0.051 | 0.229 | 0.022 |
| From State 2 | 0.037 | 0.933 | 0.027 | 0.003 |
| From State 3 | 0.231 | 0.018 | 0.560 | 0.191 |
| From State 4 | 0.024 | 0.002 | 0.222 | 0.752 |

**Supplementary Table S2.**
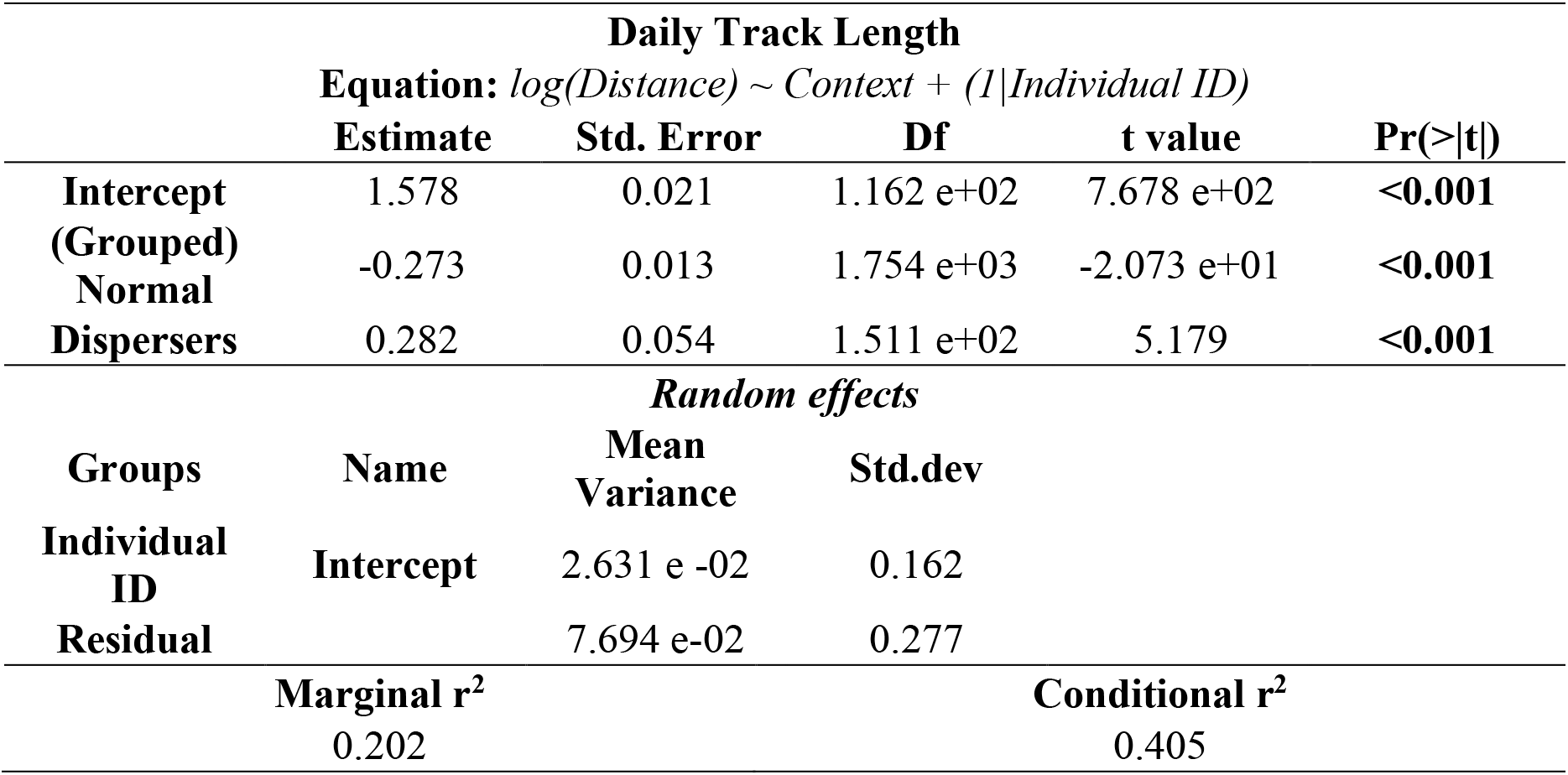
Results of the LMM for log-transformed daily track length, in kilometers, and its response to categorical context (i.e. group member during a day of large movements, group member on a normal day, or lone disperser). Reference level of contexts is group members during large movements.

**Supplementary Table S3.** Results of the LMM for total daily energy expenditure from movement, in kilojoules kg^-1^ day^-1^, and its response to categorical context (i.e. group member during a day of large movements, group member on a normal day, or lone disperser). Reference level of contexts is group members during large movements.

| Daily Energetic Expenditure |  |  |  |  |  |
| --- | --- | --- | --- | --- | --- |
| Equation: *Daily energy expenditure ~ Context + (1/Individual ID)* |  |  |  |  |  |
|  | Estimate | Std. Error | df | t value | Pr(>\|t\|) |
| Intercept | 4.789 e+02 | 2.781 | 7.806 e+01 | 1.722 e+02 | <0.001 |
| (Grouped) Normal | -1.351 e+01 | 3.065 | 4.885 e+02 | -4.408 | <0.001 |
| Dispersers | 2.860 e+01 | 6.817 | 8.735 e+01 | 4.196 | <0.001 |
| *Random effects* |  |  |  |  |  |
| Groups | Name | Mean Variance | Std.dev |  |  |
| Individual ID | Intercept | 1.683 e+02 | 12.97 |  |  |
| Residual |  | 1.169 e+03 | 34.19 |  |  |
| Marginal r<sup>2</sup> |  |  | Conditional r<sup>2</sup> |  |  |
| 0.096 |  |  | 0.210 |  |  |

**Supplementary Table S4.**
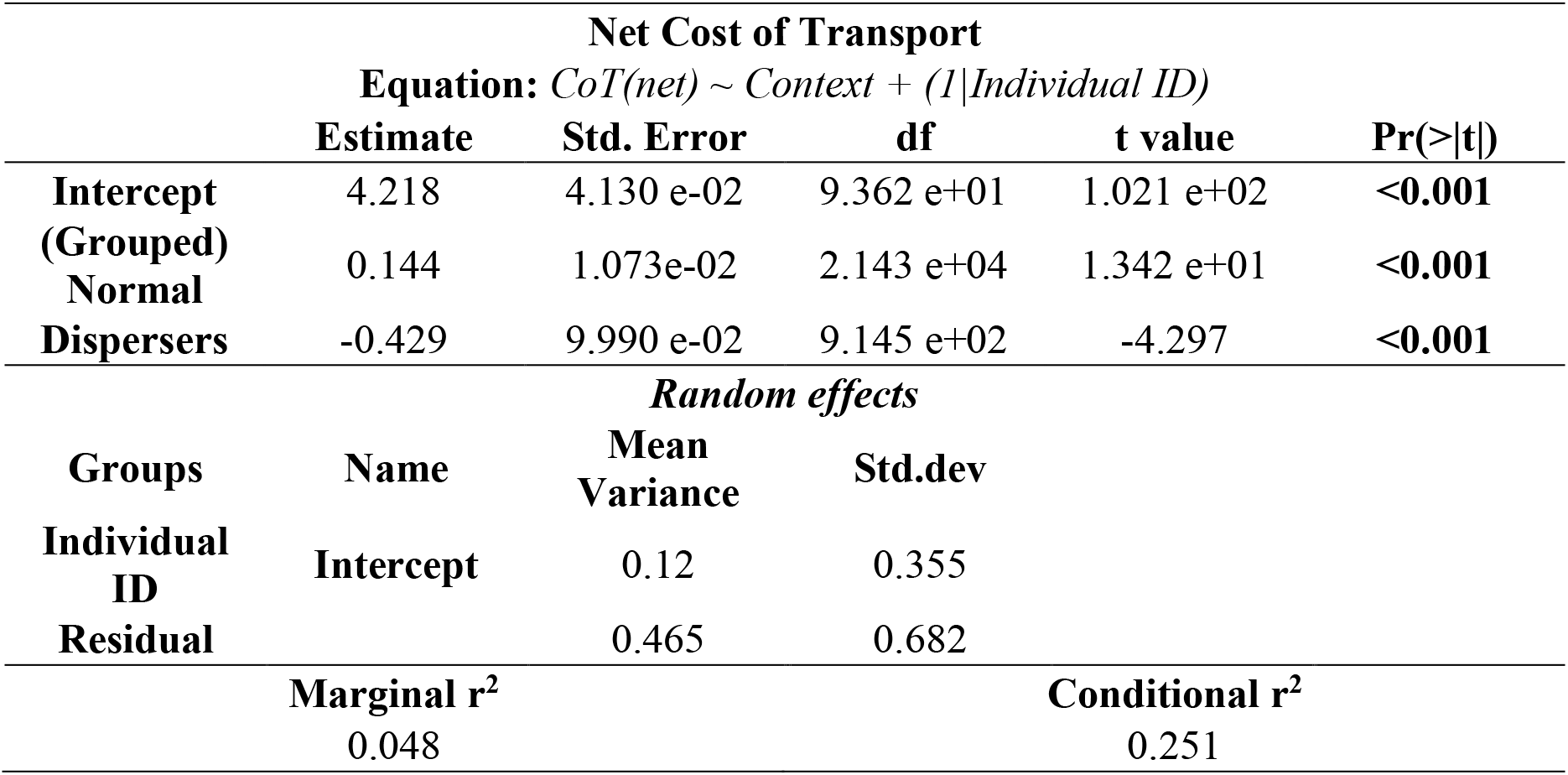
Results of the LMM for cost of transport (CoT), in Joules kg^-1^ m^-1^, over 50m of net displacement, and its response to categorical context (i.e. group member during a day of large movements, group member on a normal day, or lone disperser). Reference level of contexts is group members during large movements.

**Supplementary Table S5.**
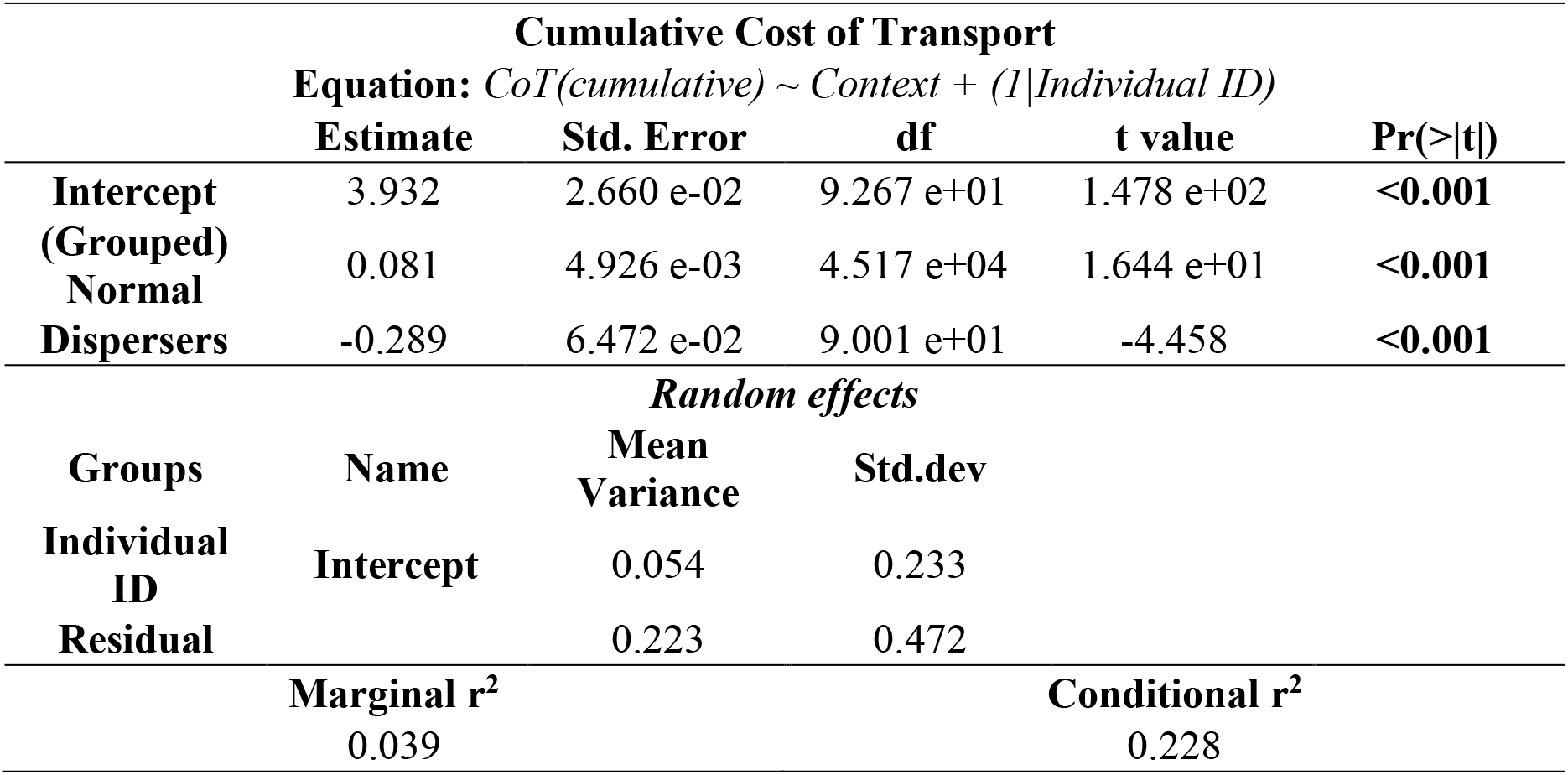
Results of the LMM for cost of transport (CoT), in Joules kg^-1^ m^-1^, over 50m of cumulative displacement, and its response to categorical context (i.e. group member during a day of large movements, group member on a normal day, or lone disperser). Reference level of contexts is group members during large movements.

**Supplementary Table S6.** Results of the LMM for movement speed (i.e. speed-while-moving) in m s^-1^, and its response to categorical context (i.e. group member during a day of large movements, group member on a normal day, or lone disperser). Reference level of contexts is group members during large movements.

| Movement Speed |  |  |  |  |  |
| --- | --- | --- | --- | --- | --- |
| Equation: *Speed~ Context + (1/Individual ID)* |  |  |  |  |  |
|  | Estimate | Std. Error | df | t value | Pr(>\|t\|) |
| Intercept | 0.281 | 8.096e-03 | 7.949 e+01 | 3.474 e+01 | <**0.001** |
| (Grouped) Normal | -0.029 | 1.696 e-03 | 4.566 e+04 | -1.696 e+01 | <**0.001** |
| Dispersers | 0.073 | 1.980 e-02 | 7.757 e+01 | 3.708 | <**0.001** |
| *Random effects* |  |  |  |  |  |
| Groups | Name | Mean Variance | Std.dev |  |  |
| Individual ID | Intercept | 5.018 e-03 | 0.071 |  |  |
| Residual |  | 2.819 e-02 | 0.168 |  |  |
| Marginal r<sup>2</sup> |  |  | Conditional r<sup>2</sup> |  |  |
| 0.022 |  |  | 0.170 |  |  |

**Supplementary Table S7.** Results of the LMM for straightness of movement (a straightness index, SI) over 5-minute intervals, and its response to categorical context (i.e. group member during a day of large movements, group member on a normal day, or lone disperser). Reference level of contexts is group members during large movements.

| Straightness of Movement |  |  |  |  |  |
| --- | --- | --- | --- | --- | --- |
| Equation: *Straightness ~ Context + (1/Individual ID)* |  |  |  |  |  |
|  | Estimate | Std. Error | df | t value | Pr(>\|t\|) |
| Intercept | 4.797 e-01 | 8.510 e-03 | 8.181 e+01 | 5.637 e+01 | <0.001 |
| (Grouped) Normal | -5.165 e-02 | 2.426 e-03 | 4.793 e+04 | -2.129 e+01 | <0.001 |
| Dispersers | -9.042 e-03 | 2.064 e-02 | 7.931 e+01 | -0.438 | 0.663 |
| *Random effects* |  |  |  |  |  |
| Groups | Name | Mean Variance | Std.dev |  |  |
| Individual ID | Intercept | 5.236 e-03 | 0.072 |  |  |
| Residual |  | 6.086 e-02 | 0.247 |  |  |
| Marginal r<sup>2</sup> |  |  | Conditional r<sup>2</sup> |  |  |
| 0.009 |  |  | 0.088 |  |  |

**Supplementary Figure S1.**
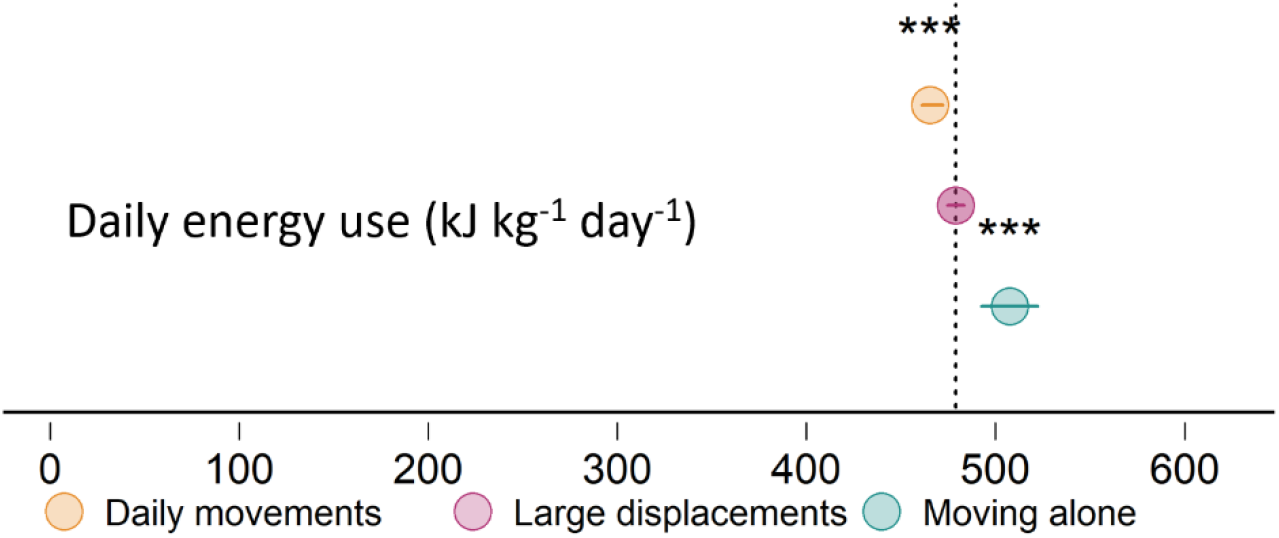
Birds moving in groups expended more energy while moving during big days of movement (purple) than they do normally (orange), but less than lone dispersers. Summary values from a linear misxed-effect model (LMMs) characterizing the energy consumed due to movement over a 13-hour day, across categories (coefficient±95% confidence intervals), with significance (*** p<0.001) estimated using big day movements of group members as the reference category (dotted vertical line corresponds to model intercept).

